# FoQDE2-dependent milRNA promotes *Fusarium oxysporum* f. sp. *cubense* virulence by targeting a glycosyl hydrolase coding gene at transcriptional level

**DOI:** 10.1101/2021.12.02.470887

**Authors:** Minhui Li, Lifei Xie, Meng Wang, Yilian Lin, Yong Zhang, Jiaqi Zhong, Jing Zeng, Guanghui Kong, Pingen Xi, Huaping Li, Li-Jun Ma, Zide Jiang

## Abstract

MicroRNAs (miRNAs) are small non-coding RNAs that regulate protein-coding gene expression primarily found in plants and animals. Fungi produce microRNA-like RNAs (milRNAs) that are structurally similar to miRNAs and functionally important in various biological processes. The fungus *Fusarium oxysporum* f. sp. *cubense* (*Foc*) is the causal agent of Panama disease that threatens global banana production. It remains uncharacterized about the biosynthesis and functions of milRNAs in *Foc*. In this study, we investigated the biological function of milRNAs contributing to *Foc* pathogenesis. Within 24 hours post infecting the host, the Argonaute coding gene *FoQDE2*, and two Dicer coding genes *FoDCL1* and *FoDCL2*, all of which are involved in milRNA biosynthesis, were significantly induced. *FoQDE2* deletion mutant exhibited decreased virulence and hypersensitivity to hydrogen peroxide (H_2_O_2_). These results indicate that milRNA biosynthesis is crucial for *Foc* pathogenesis. By small RNA sequencing, we identified 364 small RNA-producing loci in the *Foc* genome, 25 of which were significantly downregulated in the *FoQDE2* deletion mutant, from which milR-87 was verified as a FoQDE2-depedent milRNA based on qRT-PCR analysis. Through deletion and overexpression of milR-87 in the wild-type *Foc* strain, functions of milR-87 were studied. The results showed that milR-87 is crucial for *Foc* virulence in infection process. We furthermore identified a glycosyl hydrolase-coding gene, *FOIG_15013*, as the direct target of milR-87. The *FOIG_15013* deletion mutant displayed a dramatic increase in the growth, conidiation and virulence. Transient expression of FOIG_15013 in *Nicotiana benthamiana* leaves activates the host defense responses. Collectively, this study documents the involvement of milRNAs in the manifestation of the devastating fungal disease in banana, and demonstrates the importance of milRNAs in the pathogenesis and other biological processes. Further analyses of the biosynthesis and expression regulation of fungal milRNAs may offer a novel strategy to combat devastating fungal diseases.

**Author summary:** The fungus *Fusarium oxysporum* f. sp. *cubense* (*Foc*) is the causal agent of Panama disease that threatens global banana production. As a typical representative of *F. oxysporum* species complex, the pathogen has been widely concerned. However, pathogenesis of *Foc* is not fully elucidated. In particular, pathogenic regulatory mechanism of the microRNA like small RNAs (milRNAs) found in *Foc* is unknown. Here, we found that *FoQDE2,* one Argonaute coding gene, and two Dicer coding genes *FoDCL1* and *FoDCL2*, which are involved in milRNA biosynthesis, are significantly induced during the early infection stage of *Foc*. The results suggested that the milRNAs biosynthesis mediated by these genes may play an active role in the infection process of *Foc*. Based on this assumption, we subsequently found a FoQDE2-dependent milRNA (milR-87) and identified its target gene. Functional analysis showed that FoQDE2, miR-87 and its target gene were involved in the pathogenicity of *Foc* in different degree. The studies help us gain insight into the pathogenesis with FoQDE2, milR-87, and its target gene as central axis in *Foc*. The identified pathogenicity-involved milRNA provides an active target for developing novel and efficient biocontrol agents against Panama disease.

## Introduction

Banana *Fusarium* wilt, also known as Panama disease, is caused by the fungal pathogen *Fusarium oxysporum* f. sp. *cubense* (*Foc*). And it poses a serious threat to the banana industry worldwide [1]. Three physiological races of *Foc* have been identified, of which tropical race 4 (TR4) has the most devastating effect on banana production [2]. TR4 originated in Southeast Asia and is the main cause of banana *Fusarium* wilt in China [2–5]. TR4 has spread rapidly around the world, and has been reported in Mozambique, Australia, Pakistan, and even in countries along the Mediterranean coast, in countries such as Lebanon, Oman, and Jordan [1, 6]. However, fungicides, flood fallowing, and organic amendments have rarely provided long-term control in any banana planting area. The only effective method for controlling the dissemination and subsequent infections by *Foc* in banana is by the quarantine or exclusion of infected properties or by planting non-host crops or cultivars [7]. Lack of effective methods to control banana fusarium wilt seriously imperils global banana production. Improved strategies to control this devastating disease are urgently needed.

MicroRNAs (miRNAs), a type of small non-coding single-stranded RNAs, play crucial roles in diverse biological processes [8, 9]. Through base pairing with target messenger RNAs (mRNAs), miRNAs degrade the target mRNA or inhibit its translation and thereby regulate gene expression at the post transcription level [10]. Small RNAs (sRNAs) have been reported in various fungi, including *Neurospora crassa*, *Magnaporthe oryzae*, *Botrytis cinerea*, and *Sclerotinia sclerotiorum* [11, 14]. Because some of these sRNAs are structurally similar to miRNAs from plants and animals, they are called microRNA-like RNAs (milRNAs) [11, 13]. In *B. cinerea*, milRNAs have been identified as virulent effectors that suppress host immunity and facilitate fungal infection [13, 15]. In *Verticillium dahliae*, a novel milRNA, VdmilR1, was reported to play crucial roles in pathogenicity [16]. Recently, a milRNA (Vm-milR37) expressed exclusively in the mycelium, was verified to contribute to pathogenicity in *Valsa mali* [17]. On the other hand, plants also export miRNAs or siRNAs to inhibit gene expression in fungal pathogens and confer efficient crop protection from pathogen infection [18–20]. Thus the trans-kingdom sRNAs play key roles in host-pathogen interactions [21].

Four types of milRNAs generated from four different biosynthesis pathways, namely milR-1 to -4, have been reported in the model fungus *N. crassa* [11]. Different combinations of factors, including Dicers, QDE2 (Quelling Deficient 2), the exonuclease QIP (QDE2 interacting protein), and an RNAse III domain-containing protein, MRPL3, are involved in the production of miRNAs [11,22,23]. The reported milRNA biosynthesis pathways in *N. crassa* appear more complex and diverse than that in plants and animals [24].

Argonaute (AGO) proteins are evolutionarily conserved in all domains of life and play a key role in the RNA interference (RNAi) pathway [25]. As in plants and animals, AGO proteins are the core components of the RNA-induced silencing complex (RISC) and contribute to gene silencing by using the loaded sRNA as a specificity determinant in fungi [14]. QDE2, one of the two identified AGO proteins in *N. crassa*, functions as a slicer and is required for the biogenesis of some sRNAs such as milRNAs and PIWI-interacting RNAs [11, 23]. Suppressor of meiotic silencing 2 (Sms-2), another reported AGO protein in the *N. crassa* genome, is thought to function by binding to sRNAs originating from the unpaired DNA region and is required for meiotic silencing of unpaired DNA [24, 26]. In the model fungus *N. crassa*, milR-1 type of milRNAs are the most abundant milRNAs and the maturation of the milRNAs requires the AGO protein QDE-2 [11, 23]. However, whether this type of milRNAs exixt and what function they have in *Foc* remain unclear.

In this study, we identified AGO and Dicer proteins in *Foc* and examined their functions in milRNAs biosynthesis and in fungal pathogenesis by sRNA sequencing and reverse genetics. We identified a FoQDE2-dependent milRNA (milR-87), which contributes to invasive growth during the initial stage of *Foc* infection and thus affects *Foc* pathogenicity, likely by targeting the gene *FOIG_15013*. *FOIG_15013* encodes a glycosyl hydrolase, which appears as a negative regulator of *Foc* conidiation and pathogencity. Overall, our findings uncover the novel function of milRNA in *Foc* pathogenicity.

## Results

### Identification of AGO protein FoQDE2 in *Foc*

By performing an orthologous protein BLAST (Basic Local Alignment Search Tool, https://blast.ncbi.nlm.nih.gov/Blast.cgi) search, we identified two AGO proteins in *Foc*, encoded by *FoQDE*2 (gene number: FOIG_01986) and *FoAGO2* (gene number: FOIG_01246) respectively. Conserved domain prediction showed that the proteins have four domains: a variable N-terminal domain (ArgoN), a linker 1 domain (ArgoL1), a PAZ domain, and a PIWI domain (Fig 1A).

**Fig 1.**
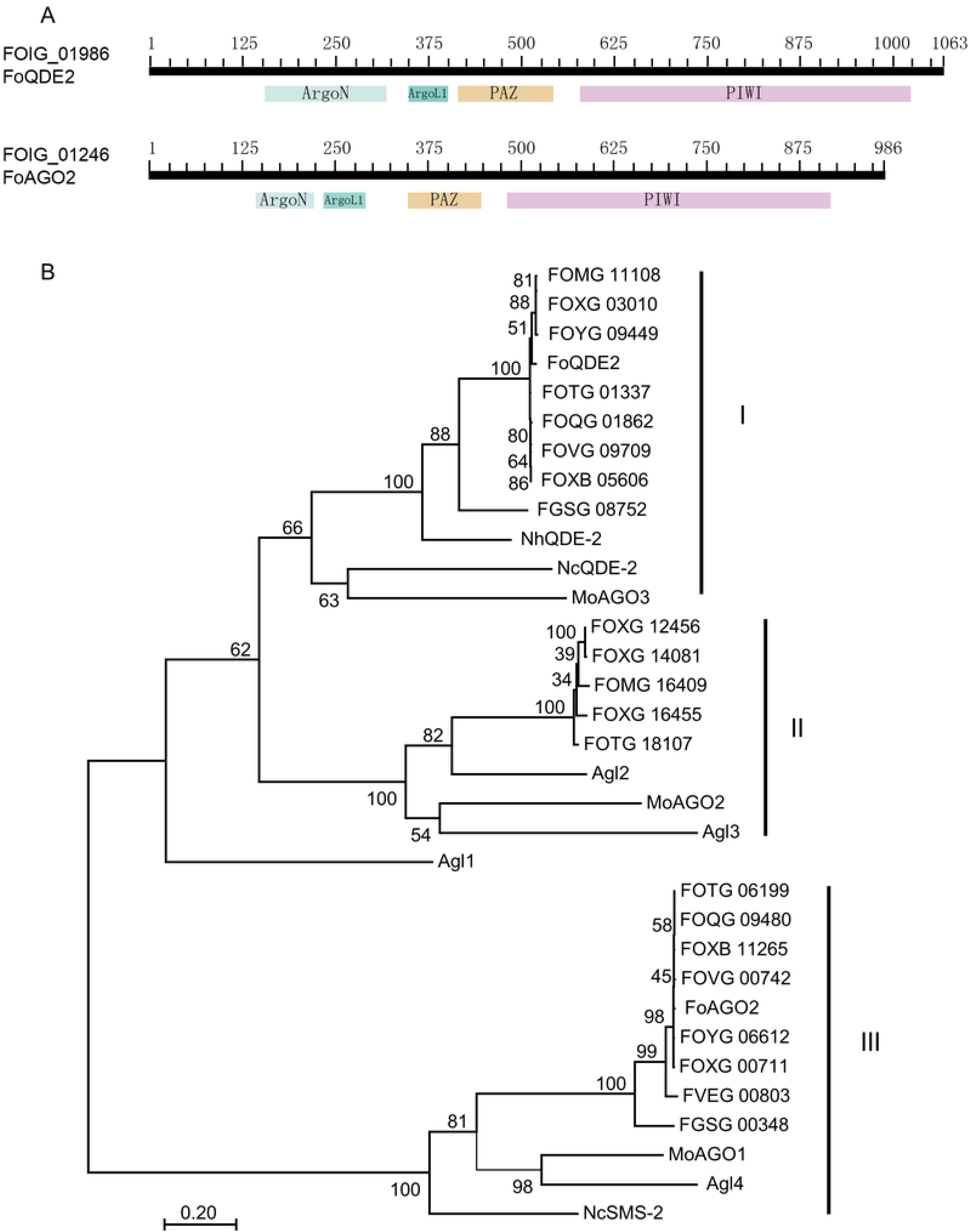
Phylogenetic analysis of fungal Argonaute proteins. (**A**) Conserved domains of Argonaute proteins were predicted through BLASTp on NCBI. (**B**) Fungal Argonaute protein sequences were first aligned using Clustal×1.8 and then the aligned sequences were analyzed by the Maximum Likelihood method implemented in MEGA 7. Bootstrap values are expressed as a percentage of 1000 replicates. In addition to FOIG genes from the *Fusarium oxysporum* f. sp. *cubense* (*Foc*) genome, genes used for this analysis include Agl1–Agl4 from *Cryphonectria parasitica* (Sun et al., 2009), a chestnut blight fungus; MoAGO1–MoAGO3 from *Magnaporthe oryzae*, a rice blast fungus (Nguyen et al., 2019); NcQDE2 and NcSMS-2 from the model fungus *Neurospora crassa* (Dang et al., 2014); and genes from the *F. graminearum* (FGSG), *F. verticillioides* (FVEG), and *F. oxysporum* (FOVG: GCA_000260075.2; FOQG: GCA_000260235.2; FOTG: GCA_000260175.2; FOMG: GCA_000260495.2; FOXG: GCA_000149955.2; and FOYG: GCA_000271745.2) genomes.

The amino acid sequences of AGO orthologs in other fungal species were retrieved from GenBank for phylogenetic analysis. Two to five AGO-like proteins were identified in different strains of *F. oxysporum*, but only two in *Foc*. Phylogenetic analysis indicated that AGO proteins from filamentous fungi could be divided into three subgroups (Fig 1B). In the first group, *FoQDE2* and its orthologs from *F. oxysporum* and *F. graminearum* were clustered with QDE2 from *N. crassa*, AGO-like protein MoAGO3 from *M. oryzae* [14], and Agl1 from the chestnut blight fungus *Cryphonectria parasitica* [27]. The second group includes AGO-like proteins from *F. oxysporum* and Agl2 and Agl3 from *C. parasitica*. The third group is composed of FoAGO2 and its orthologs from *F. oxysporum* and *F. graminearum*, Agl4 from *C. parasitica*, MoAGO1 from *M. oryzae*, and SMS2 from *N. crassa* [24]. The phylogenetic analysis showed that the AGO proteins in *Foc* are orthologous to those of the other filamentous fungi.

### Characterization of sRNA biosynthesis-related genes in *Foc*

In addition to the AGO coding genes, two Dicer coding genes (*FoDCL1* and *FoDCL2*) exist in *Foc*, which function in the biosynthesis of fungal sRNAs [11]. Compared with *Foc* in pure culture conditions, *FoQDE2* and Dicer encoding genes were significantly upregulated at 24 hours post inoculation (hpi) on host banana plants. In contrast, transcription of *FoAGO2* was nearly undetectable (Fig 2B). The transcriptional induction of these sRNA biosynthesis genes during host infection stage suggests that the sRNAs synthesis mediated by these genes may play an active role during *Foc* pathogenesis.

**Fig 2.**
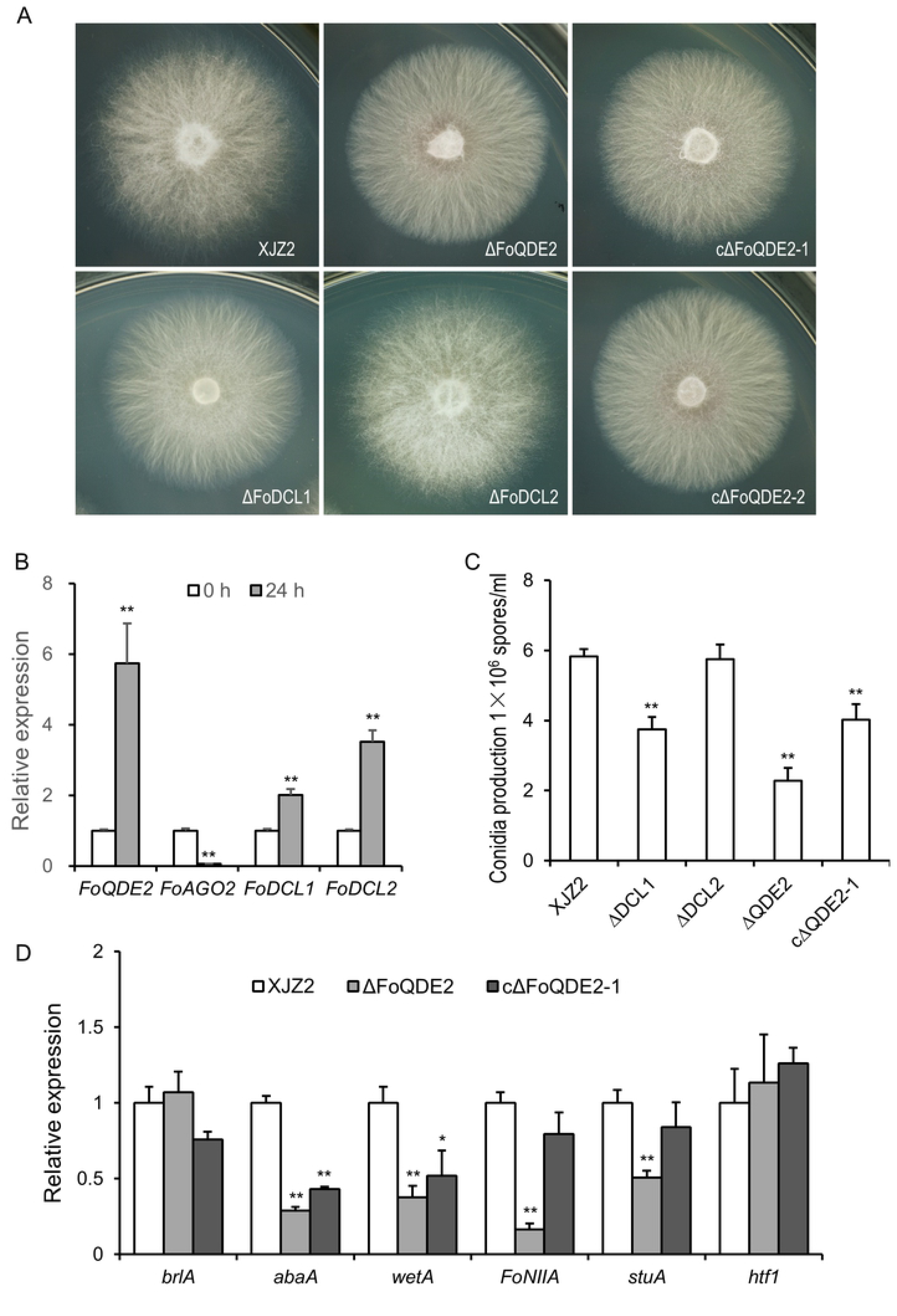
Characterization of milRNA biosynthetic pathway genes in *F. oxysporum* f. sp. *cubense*. (**A**) Colony morphology of the tested strains grown on PDA. Photographs were taken after 3 days of culture on PDA plates. (**B**) Expression patterns of milRNA biosynthetic pathway genes in pure culture conditions (0 h) and at 24 hours post inoculation (24 hpi). Relative folds were calculated by 2^-ΔΔCt^ method (Li et al., 2014), using transcription elongation factor 1 α gene (EF1α) as internal control. Mean ± S.D. was derived from three biological repeats. **, p<0.01. (**C**) Conidial production of the tested strains. The tested strains were grown on PDA plates at 28 °C for 7 days. Then same amount of colony discs were washed by water supplemented with 0.05% tween 20 to prepare conidia suspension. And conidia were quantified under microscope using a haemocytometer. Error bars indicate S. D. (n=10). **, p<0.01. (**D**) Expression patterns of the conidial production-related genes in the wild type strain XJZ2, the Δ*FoQDE2* mutant and complimentary strain cΔ*FoQDE2*-1. RNAs were isolated from freshly cultured mycelia and were reverse transcribed into cDNAs. Transcript levels of the candidate genes in the Δ*FoQDE2* mutant and complimentary strain cΔ*FoQDE2*-1 are presented as values relative to the expression in the wild-type strain XJZ2. Error bars indicate S. D. (n=3). **, p<0.01.

To further investigate the function of the three upregulated genes (*FoQDE2*, *FoDCL1*, and *FoDCL2*), we generated these genes deletion mutants in the wild-type (WT) strain XJZ2 via homologous recombination (S1A Fig). Four independent *FoQDE2-*deletion mutants were verified by PCR and Southern blot analysis (S1B and S1C Fig). Four *FoDCL1-*deletion mutants and *FoDCL2-*deletion mutants were respectively obtained and verified by Southern blot analysis (S1C Fig). Four complemented strains were also generated by expressing the *FoQDE2* locus (S1A Fig) in one of the *ΔFoQDE2* mutant, as confirmed by PCR (S1D Fig) with primer pairs FoQDE2-F1/FoQDE2-R, HYG-F/HYG-R2, and Zeo-F/Zeo-R (position of the primer pairs were presented in S1A Fig; sequences listed in S1 Table).

Relative transcript levels of *FoQDE2*, *FoDCL1*, and *FoDCL2* were examined by quantitative real-time PCR (qRT-PCR) analysis in the WT strain and the mutants. No *FoQDE2*, *FoDCL1*, or *FoDCL2* transcript was detected in the corresponding deletion mutants (S2A Fig), confirming that the genes had been successfully deleted in the respective mutant. Transcript level of *FoQDE2* was detected in the complemented strains, to a comparable level of that in the WT (S2B Fig), indicating that *FoQDE2* was restored in the complemented strains (cΔ*FoQDE2*-1/-2).

### FoQDE2 is required for proper mycelial growth, and conidial production in *Foc*

We next assessed the biological function of FoQDE2 by examining growth and conidiation phenotypes in the Δ*FoQDE2* mutants. The results showed that, the colonial morphology of the Δ*FoQDE2* was strikingly different from that of the WT strain XJZ2. The aerial hyphae of Δ*FoQDE2* were much less abundant than those of the WT, and the mycelia grew close to the surface of the PDA media. Such morphological change of the colonies was restored to different degrees in the two complemented *FoQDE*2 strains (cΔ*FoQDE2*-1 and cΔ*FoQDE2*-2; Fig 2A). On the other hand, the colony morphology was not much changed in the Δ*FoDCL1* or Δ*FoDCL2* mutants (Fig 2A).

We assessed mycelial growth rate in the WT and mutant strains. From the fourth day of culture, the growth of Δ*FoQDE2* was significantly slower than that of the WT (*p*<0.01). The growth of the complemented strains was not fully recovered. The Δ*FoDCL1* and Δ*FoDCL2* mutants displayed no difference in mycelial growth compared with the WT (S2 Table). Thus, loss of *FoQDE2* led to slow mycelial growth on PDA medium.

Furthermore, microconidia production was significantly reduced in the Δ*FoQDE2* mutant, compared to the WT, when cultured on PDA medium for 7 days. Such conidiation defect could be partially restored in the *FoQDE2* complemented strains (Fig 2C). Compared with the WT, Δ*FoDCL1* also produced less microconidia, whereas Δ*FoDCL2* showed no significant difference in conidial production (Fig 2C).

To elucidate the causes of the decreased conidia production in Δ*FoQDE2*, six reported conidiation-related genes [28–31] were selected for their transcript levels assessment by qRT-PCR. Compared with the WT, the transcript levels of four selected conidiation-related genes, *StuA*, *FoNIIA*, *AbaA*, and *WetA*, were significantly down-regulated in Δ*FoQDE*2. Transcription of *StuA* and *FoNIIA* were restored in the *FoQDE2* complemented strain (cΔ*FoQDE2*-1), to a level comparable to the WT (Fig 2D). The transcript levels of *brlA* and *Htf1* were not affected by the loss of *FoQDE2* (Fig 2D). Overall, our results showed that deletion of *FoQDE2* leads to morphological changes, including changes in colonial morphology and reduction of mycelial growth, as well as reduction in conidial production of *Foc*.

### FoQDE2 is required for oxidative stress tolerance

We next tested the sensitivities of the WT and the mutant strains to oxidative stress. The Δ*FoQDE2* mutant was hypersensitive to 3mM H_2_O_2_ when cultured on minimal medium (MM), which could be restored in the complemented strains (Fig 3A and 3B). However, the Δ*FoDCL1* and Δ*FoDCL2* mutants showed no difference in sensitivity to oxidative stress as compared to the WT (Fig 3A and 3B). Taken together, FoQDE2 is required for tolerance to oxidative stress.

**Fig 3.**
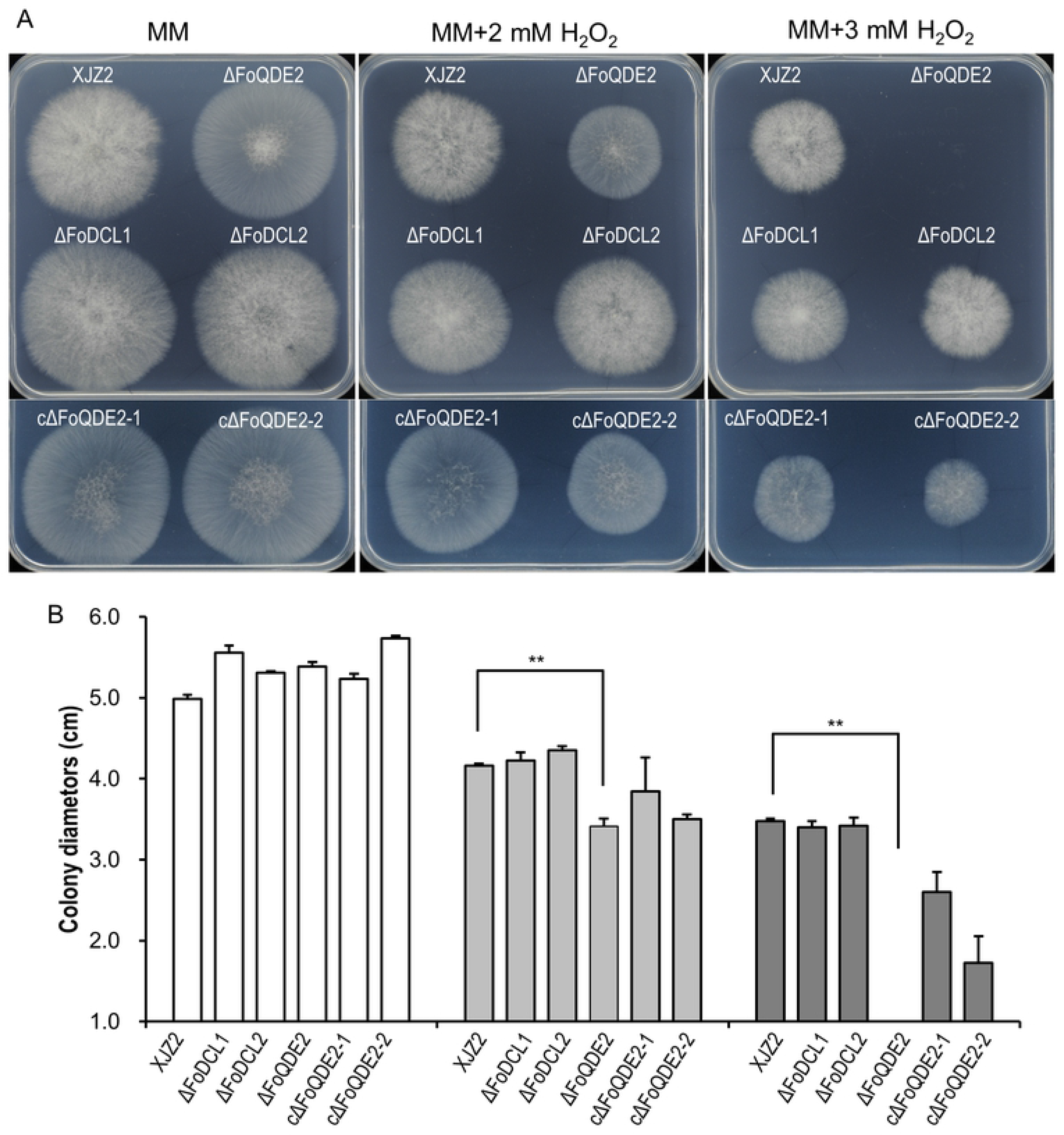
Mycelial sensitivity to H_2_O_2_. (**A**) Mycelial growth of milRNA biosynthetic pathway gene knockout mutants (Δ*FoQDE2*, Δ*FoDCL1*, and Δ*FoDCL2*) and two *FoQDE2* complementation transformants (cΔ*FoQDE2*-1 and cΔ*FoQDE2*-2) under oxidative stress. The wild type (WT) strain XJZ2, mutants Δ*FoQDE2*, ΔFoDCL1, and ΔFoDCL2, and the cΔ*FoQDE2*-1 and cΔ*FoQDE2*-2 were inoculated on minimal medium (MM) with 0, 2, or 3 mM H_2_O_2_ and cultured at 28°C for 4 days. (**B**) Measurements and statistical analysis of the colony diameters of the tested strains. A *t*-test was used to identify significant differences between the WT and the mutants. Error bars indicate S. D. (n=3). **, p<0.01.

### FoQDE2 is involved in the virulence to banana seedlings

Mycelial infection capacity was tested on the tomato fruit surface and the incidence of tomato fruit infection was recorded at five days post inoculation (dpi). The Δ*FoQDE2* mutant did not penetrate the epidermis or cause fruit tissue necrosis on the surface of tomato as the WT and complemented strains did (Fig 4A). In contrast, the Δ*FoDCL1* and Δ*FoDCL2* mutants caused tissue necrosis similar to that of WT (Fig 4A). Thus, FoQDE2 is required for successful invasive growth.

**Fig 4.**
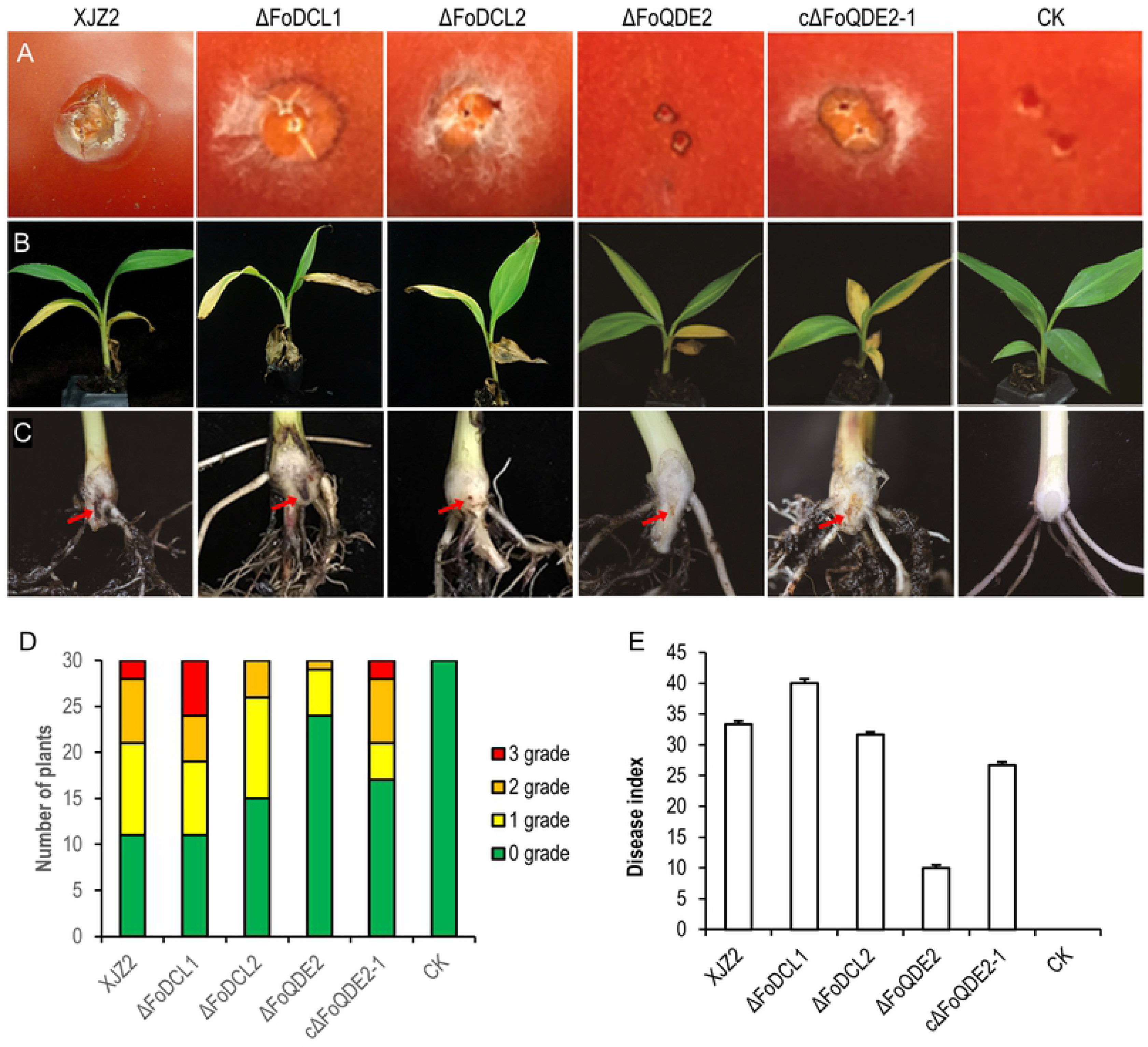
Pathogenicity assay. (**A**) Invasive growth on tomato fruits. The surfaces of tomato fruits were inoculated with the WT strain XJZ2, Δ*FoQDE2* mutant, the cΔ*FoQDE2*-1 complementation transformant, and water control (CK). The incidence of necrosis on tomato fruits was assessed at 5 days post inoculation. (**B**) and (**C**) Symptoms of pathogenicity on the leaves (B) and corm (C) of the banana plant. (**D**) and (**E**) Disease severity analyzed by diseased plantlet number of different disease grade (D) and disease index (E). The disease grade was classified as 0 (no symptoms in the corm of the banana plantlet), 1 (the presence of small vascular discoloration like brown dots), 2 (up to 50% of vascular discoloration) and 3 (greater than 50% of vascular discoloration).

Pathogenicity tests on banana (Cavendish) seedlings showed that the Δ*FoQDE2* mutant was unable to produce obvious vascular discoloration in the corm of banana seedling (Fig 4B and 4C) and was significantly reduced in virulence on banana compared to the WT (*p*<0.01, Fig 4D and 4E). The disease index of the complemented strain (cΔ*FoQDE2*-1) was increased but did not reach the level of the WT (Fig 4E), which exhibited brown discoloration in the corm of banana seedlings. On the other hand, pathogenicity of the Δ*FoDCL1* and Δ*FoDCL2* mutants were similar to that of WT (Fig 4A-4E). These observations indicate that *FoQDE2* contributes to the virulence of *Foc*.

### Identification of FoQDE2-dependent milRNA in *Foc*

To investigate the function of FoQDE2 in sRNA production, we sequenced sRNAs from the WT strain and Δ*FoQDE2* mutant. The sequencing data were deposited in the NCBI sequence read archive (SRA) under accession number PRJNA562097. Origin analysis of the sRNAs showed that reads mapped to tRNA in Δ*FoQDE2* was significantly reduced by approximately 40% than in the WT, whereas significantly more reads mapped to the UTRs, intronic, and intragenic regions in Δ*FoQDE2* than in the WT (Fig 5A). sRNA length distribution analysis showed that most reads with length of between 16–24 nt were in both WT and Δ*FoQDE2* mutant, while reads of 21 and 22 nt were reduced in the Δ*FoQDE2* mutant compared to the WT (Fig 5B). Reads starting with A or U were more abundant than those starting with C or G (Fig 5C). To identify sRNA-producing loci, reads with counts of 10 or higher that mapped to the UTR and intronic and intergenic regions were less than 300 nt were considered to be small RNA-producing loci. Using this method, 364 loci were captured in the WT and Δ*FoQDE2* mutant. Among them, 25 loci were significantly decreased and 13 loci were significantly increased in Δ*FoQDE2* compared with the WT. (Fig 5D).

**Fig 5.**
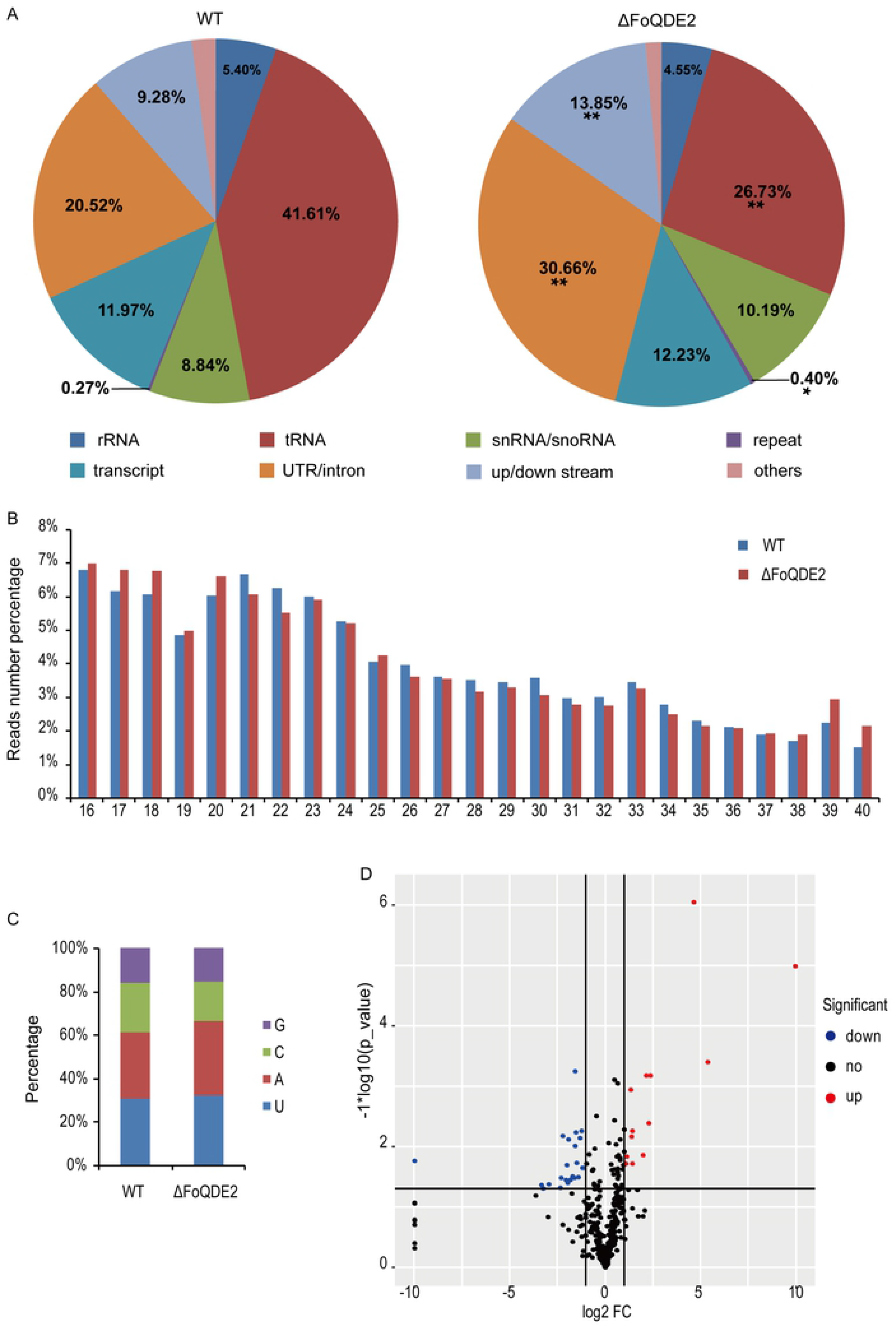
Small RNA sequencing analysis in *F. oxysporum* f. sp. *cubense*. (**A**) Characterization of small RNA-producing loci. (**B**) Length distribution of small RNAs. (**C**) Analysis of the 5′ end nucleotide preference of small RNAs. (**D**) Differential analysis of small RNA expression in the WT and *FoQDE2* deletion mutant.

To identify FoQDE2-dependent milRNA, sRNA read counts obtained from the WT and Δ*FoQDE2* were normalized and compared. To eliminate interference from sRNAs produced by mRNA degradation, the reads mapped to ORFs were discarded. A total of 25 sRNA-producing loci were significantly decreased in Δ*FoQDE2* compared with the WT. Sequences of the loci were extracted from the *Foc* II5 genome and uploaded to the RNAfold web server (http://rna.tbi.univie.ac.at//cgi-bin/RNAWebSuite/RNAfold.cgi) for RNA advanced structure prediction. The loci with a stem and loop RNA structure were validated as precursors of milRNAs. Among them, we focused on one sRNA (milRNA-87) that formed a stem and loop precursor (Fig 6A). The mature milR-87 was predicted to be of 23 nt (Fig 6A). We further verified expression level of milR-87 using qRT-PCR (Fig 6B) and sRNA Northern blot (S3E Fig). Compared to the WT, the production of milR-87 was significantly reduced in the Δ*FoQDE2* and Δ*FoDCL2* mutants (Fig 6B and S3E Fig). Based on our sRNA analysis, milR-87 is dependent on FoQDE2 and FoDCL2 in *Foc*.

**Fig 6.**
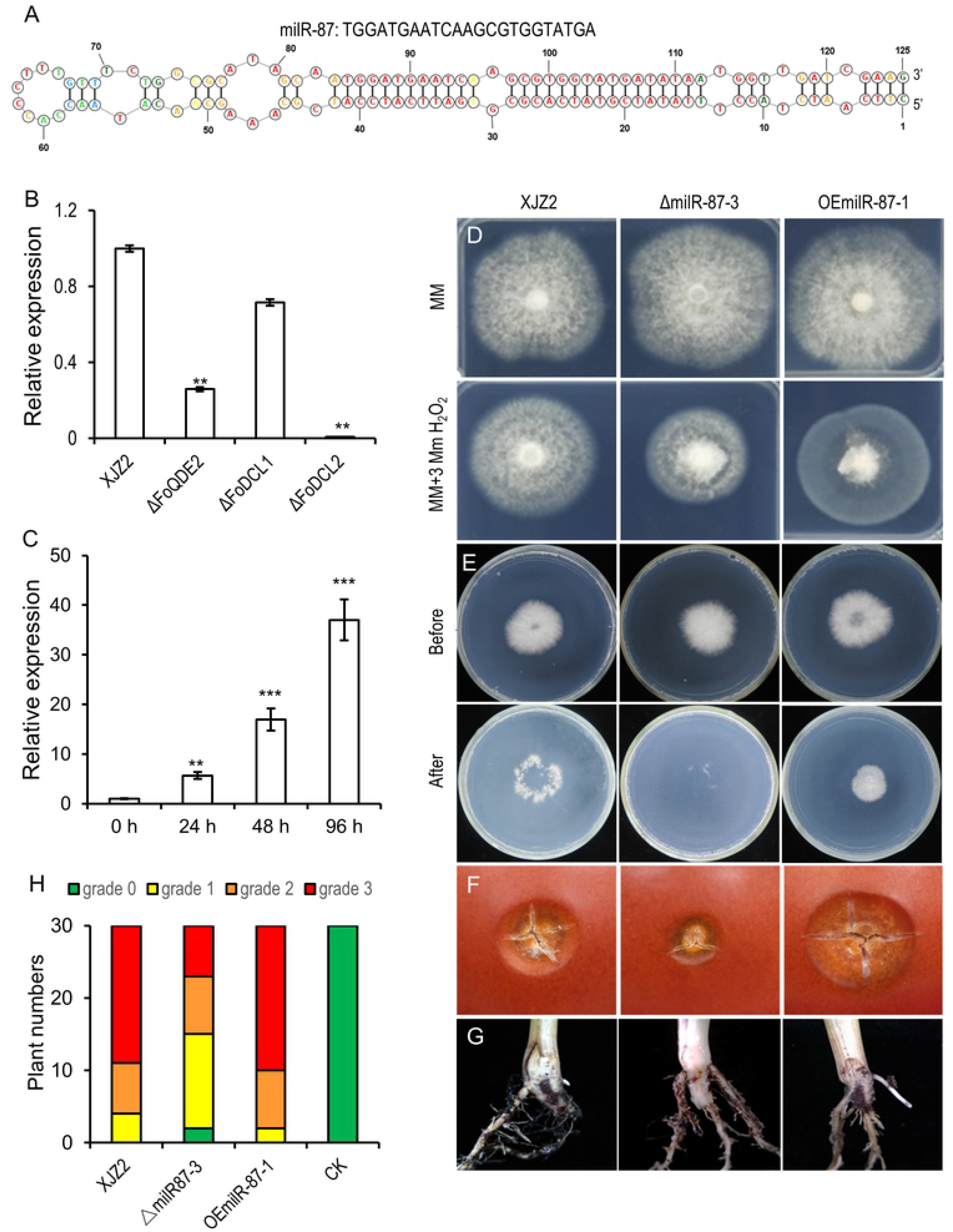
Characterization of FoQDE2-dependent milRNA (milR-87) in *F. oxysporum* f. sp. *cubense* (*Foc*). (**A**) Secondary structure of milR-87 precursor in *Foc* predicted by RNAstructure 6.3. (**B**) MilR-87 expression in different milRNA biosynthetic pathway gene deletion mutants (Δ*FoQDE2*, ΔFoDCL1, and ΔFoDCL2). (**C**) Expression patterns of milR-87 at 0, 24, 48, and 96 hours post inoculation. Total RNAs were extracted from the host banana roots inoculated with *Foc* at the different time point (0-96 h). Relative expression levels of milR-87 were calculated by 2^-ΔΔCt^ method using snRNA (U4) as internal control. Error bars indicate S. D. (n=3). **, p<0.01; ***, p<0.001. (**D**) Mycelial sensitivity to H_2_O_2_. Mycelial growth of the WT strain XJZ2, the ΔmilR-87 mutant and OEmilR-87 strain were measured at MM with 3 mM H_2_O_2_. (**E**) Cellophane penetration assay comparing the invasive growth of the XJZ2, ΔmilR-87 mutant and OEmilR-87 strain. Conidia suspensions (100 μl per strain) with same concentration of 1 × 10^5^ spores/ml were put on cellophane-covered PDA plates and incubated at 28°C for 4 days (Before, before cellophane removed), then the cellophane sheets were removed, and samples were incubated for an additional 3 days (After, after cellophane removed) and the colony size indicated mycelial growth were photographed. (**F**) Invasive growth on tomato fruits. (**G**) Symptoms of pathogenicity on the corm of the host banana seedling. (**H**) Disease severity analyzed by diseased plantlet number of different disease grade.

### milRNA (milR-87) regulates tolerance to hydrogen peroxide and is involved in the virulence of *Foc*

To determine the function of the milR-87, we generated deletion or overexpression mutants of milR-87 precursor in the WT *Foc* strain (S3A Fig). PCR results showed that the milR-87 precursor was successfully deleted or overexpressed in *Foc* (S3B and S3C Figs). Northern blot and qRT-PCR results confirmed that the milR-87 was deleted or overexpressed in the corresponding mutants (S3D and S3E Figs).

The ΔmiR-87 mutant was hypersensitive to the oxidative stress generated by 3 mM H_2_O_2_, a phenotype similar to that of the Δ*FoQDE2* mutant (Fig 6D). In contrast, the growth of milR-87 overexpress transformant (OEmilR-87-1) was not affected by oxidative stress (Fig 6D). The result indicates milR-87 positively regulates tolerance to oxidative stress in *Foc*.

Pathogenicity tests showed that the ΔmiR-87 mutant was compromised in penetrating the cellophane membrane and caused slighter necrosis on the surface of tomato fruits than the WT. While overexpressed transformant showed the opposite (Fig 6E and 6F). Infection assay using banana seedlings showed that ΔmilR-87 significantly reduced in virulence compared to the WT, while the virulence of overexpressed transformant was enhanced, as it caused obvious internal disease symptoms of brown discoloration (Fig 6G and 6H). Furthermore, qRT-PCR analysis showed that milR-87 was significantly upregulated during the early stages of infection (24, 48, and 96 hpi) compared with levels in pure culture (0 hpi) conditions (Fig 6C). These results suggest that milR-87 contributes to the infective growth and virulence of *Foc*.

To confirm the function of milRNA (milR-87) in *Foc*, synthetic double-strand siRNAs (siRNA-1 and siRNA-2) and single-strand antisense small RNA (inhibitor) were designed (S1 Table) to silence the expression of milR-87. We observed that when the siRNAs or inhibitor was transfected into *Foc* protoplast inoculated on the banana leaf, the size of the lesion decreased significantly compared to those caused by the *Foc* protoplasts with water as control (CK) (Fig 7A and 7B). An unrelated single-strand sRNA served as the negative control (NC) did not interfere with lesion formation when added into the *Foc* protoplasts inoculated on the banana leaf (Fig 7A and 7B). We noticed that the ΔmilR-87 mutant produced the smallest lesion on the banana leaf (Fig 7A and 7B). The qRT-PCR results confirmed that expression of milR-87 could be silenced by the siRNA-1, siRNA-2 and inhibitor in different degree (Fig 7C). Overall, these results confirmed that milR-87 is required for full virulence of *Foc*.

**Fig 7.**
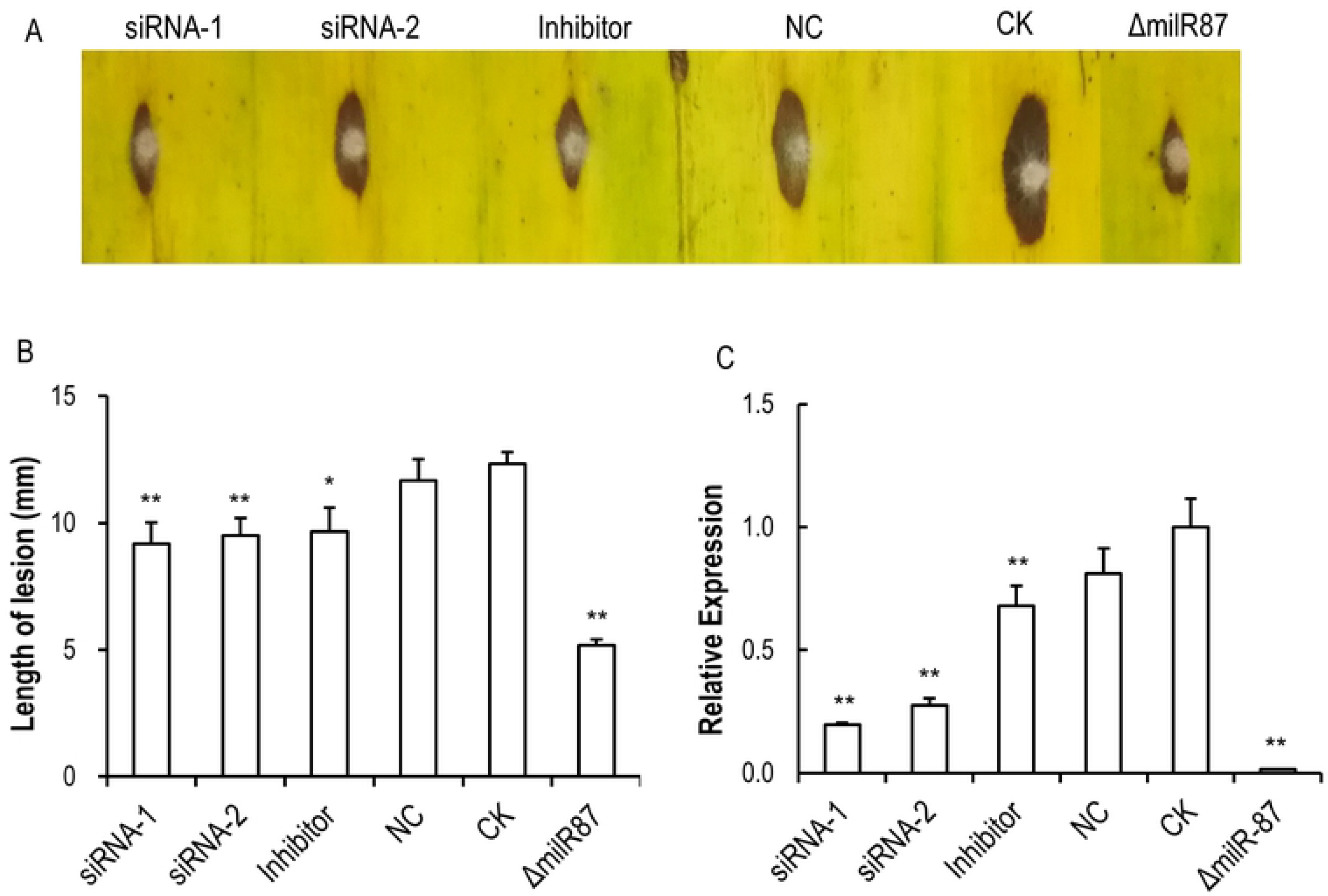
Exogenous siRNAs and inhibitor against milR-87 attenuated the virulence of *F. oxysporum* f. sp. *cubense* to banana leaves. Double-strand siRNAs (siRNA-1 and siRNA-2) and single-strand inhibitor against the precursor sequence of milR-87, as well as a random small RNA (24 nt) serving as a negative control (NC) were synthesized and transfected into *Foc* with aid of lipofectamine 2000 (Invitrogen). After 2 days transfection, conidia suspensions of the transfected strains and positive control of the ΔmilR-87 mutant, as well as non-transfected strain of *Foc* (CK) were inoculated on the banana leaves. After 3 days, the lesions on banana leaves were photographed. (**A**) Symptoms of pathogenicity on the host banana leaves. (**B**) Lesions sizes were measured for accessing virulence of the different strains. Error bars indicate S. D. (n=3). *, p<0.05; **, p<0.01. (**C**) Relative expression of milR-87 in the transfected strains, the non-transfected strain of *Foc* (CK), and the ΔmilR-87 mutant. The total RNAs of the tested strains cultured at 28°C for 7 days were extracted for milRNA detection by qRT-PCR. Relative expression levels of milR-87 in the transfected strains were normalized to that of non-transfected strain of *Foc* (CK), which was set as 1. snRNA U4 was used as internal control. Error bars indicate S. D. (n=3). **, p<0.01.

### MilRNA (milR-87) regulates the virulence by targeting a glycosyl hydrolase genes (*FOIG_15013*) at transcriptional level during the pathogenesis of *Foc*

To investigate regulatory mechanism of milR-87 in *Foc* pathogenesis, target genes were computationally predicted, and verified by qRT-PCR using total RNAs fromthe mutants (Δ*milR-87* and Δ*FoQDE2*) and WT. Totally nine genes were significantly up-regulated in the two milR-87 mutants compared to the WT (S4 Fig). One of these genes, *FOIG_15013*, encodes a glycosyl hydrolase, consisting of a signal peptide and a conserved domain of GH-79C (Fig 8A), was selected for further investigation. The predicted targeted site of milR-87 was located in the ORF region of FOIG_15013, we then generated a fusion protein by directly ligating the GFP coding sequence to the C’-terminus of FOIG_15013 (Fig 8A). A point-mutated FOIG_15013 that could not be paired with milR-87, FOIG_15013m, was also fused with GFP (Fig 8A). The constructs were respectively transformed into the Δ*miR-87* or the WT strain. This FOIG_15013-GFP fusion construct was transformed in the WT or the Δ*milR-87* mutant, to verify the possible milR-87 regulation of FOIG_15013 expression by evaluating the GFP signal brightness. In the Δ*milR-87* mutant, FOIG_15013-GFP expression was not inhibited, as indicated by the strong GFP signal (Fig 8B and 8C). In contrast, the GFP signal was hardly detected in the WT strain, likely due to milR-87 mediated suppression of FOIG_15013-GFP expression (Fig 8B and 8C). The FOIG_15013m-GFP expression was not affected in the WT background, as milR-87 was unable to pair with the mutation site. Correspondingly, transcript level of *FOIG_15013* and its mutated version in different strains was consistent with that of GFP signal detection (Fig 8D), further confirming that *FOIG_15013* is targeted by milR-87 in *Foc*.

**Fig 8.**
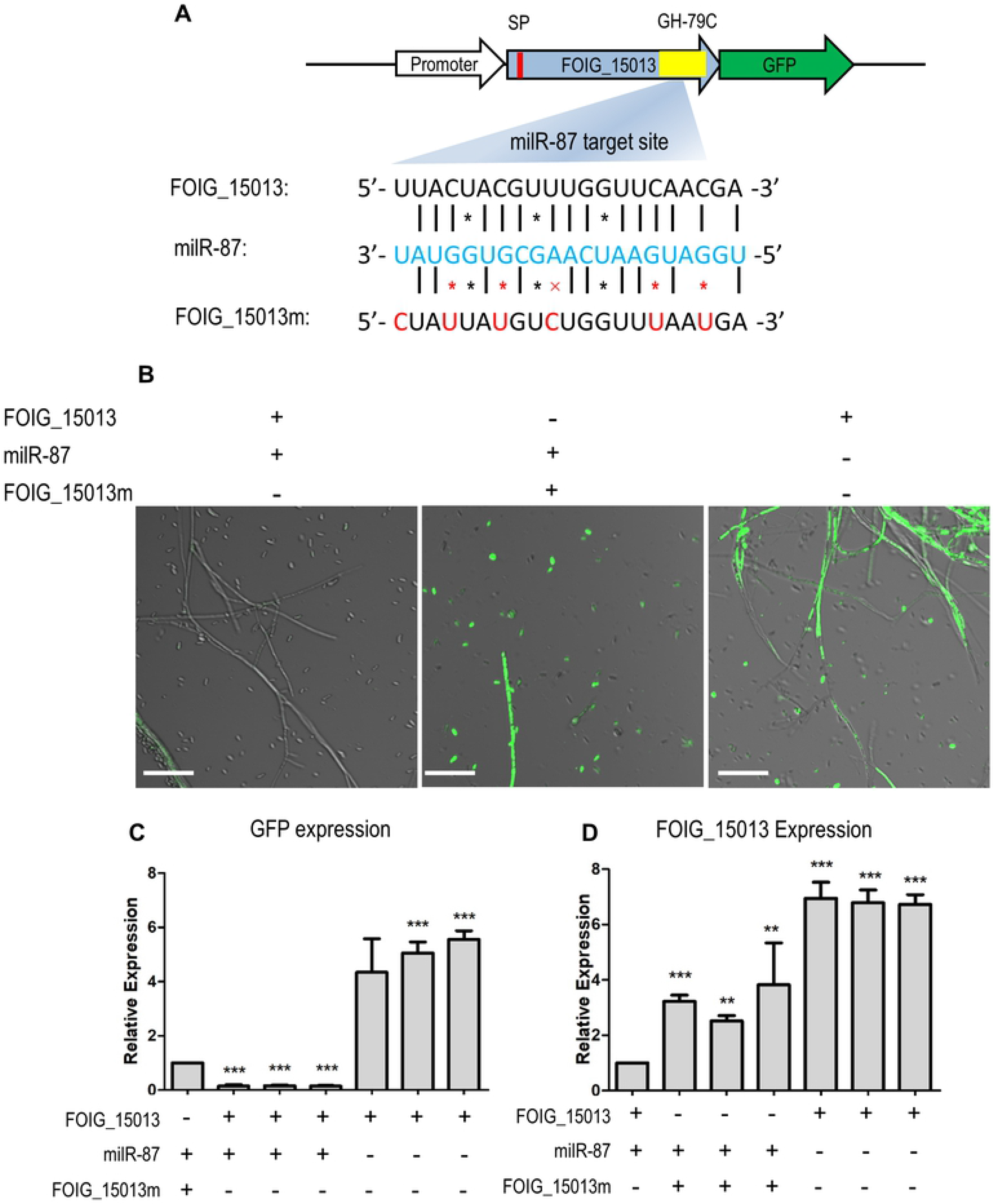
Identification of milR-87 target gene in *F. oxysporum* f. sp. *cubense*. (**A**) Schematic diagram for construct of FOIG_15013-GFP, with the wild-type or mutated (FOIG_15013m) milR-87 targeted site. The red box represents signal peptide (SP) and the yellow box represents a conserved domain of glycosyl hydrolase 79C (GH-79C) predicted in FOIG_15013. The milR-87 and its potential target sequence of FOIG_15013 was labelled in blue and black respectively. Point-mutations were in red font. (**B**) Confocal fluorescence microscopic images of the strains expressing FOIG_15013-GFP or FOIG_15013m-GFP, with or without milR-87. Bar=50 μm. (**C**) Transcript level of marker gene GFP in the tested strains. GFP expression values in different strains were normalized to that of point-mutation strain, which GFP value was set as 1. (**D**) Transcript level of FOIG_15013 in different strains. FOIG_15013 expression values in different strains were normalized to that of the GFP-labelled WT strain. The transcription elongation factor 1 α gene (EF1α) was used as internal control. Error bars indicate S.D. (n=3). **, p<0.01; ***, p<0.001.

In order to clarify whether milR-87 regulates *Foc* pathogenesis through modulating *FOIG_15013* expression, we generated the Δ*FOIG_15013* mutants (S5A and S5B Figs) and characterized them in growth and pathogenicity. The results showed that, the Δ*FOIG_15013* mutants grew faster and produced significantly more conidia than the WT strain XJZ2 when cultured on PDA plate (Fig 9A and 9B). Pathogenicity tests showed that the Δ*FOIG_15013* mutants were more virulent than the WT (Fig 9C and 9D). These results indicate that milR-87 may promote *Foc* pathogenicity via suppression of *FOIG_15013* expression, which is a negative regulator of *Foc* mycelial growth and virulence.

**Fig 9.**
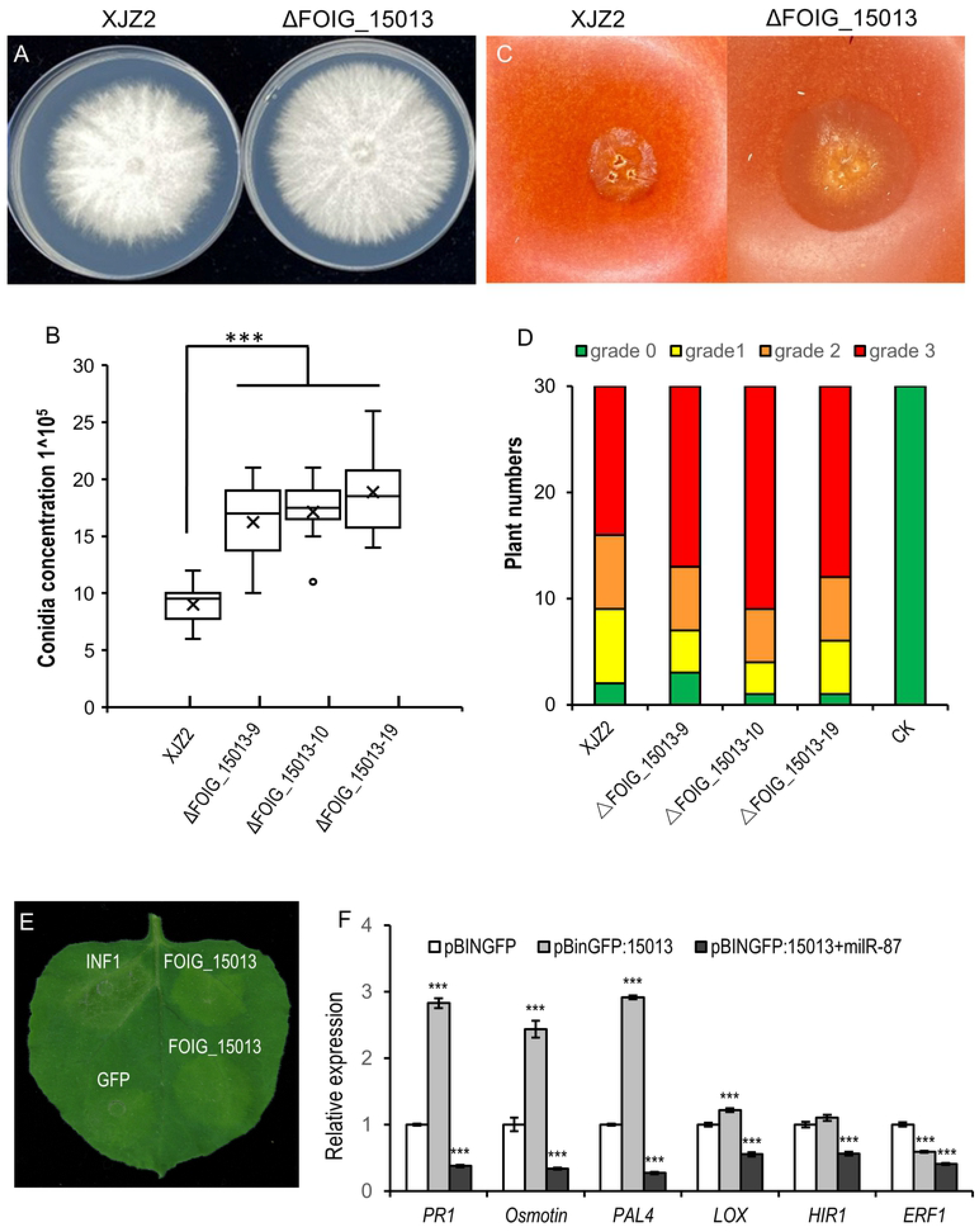
Investigation of FOIG_15013 function in *Foc* growth, conidiation, and pathogencity. (**A**) Colony morphology of the wild type strain XJZ2 and *FOIG_15013* deletion mutant (Δ*FOIG_15013*). (**B**) Conidial production of the tested strains. (**C**) Symptom of necrosis caused by the WT and the ΔFOIG_15013 mutant on the surface of tomato fruits. (**D**) Statistical results of disease severity analyzed by diseased host banana seedling number of different disease grade. (**E**) Hypersensitive response induced by transient expression of *FOIG_15013* in *N. benthamiana* leaves. Leaves of *N. benthamiana* were infiltrated with *Agrobacterium tumefaciens* carrying pBIN::GFP- *FOIG_15013* (FOIG_15013). Photographs were taken at 2 days post-agroinfiltration. INF1 and GFP and were used as positive and negative control. (**F**) Transcript levels of defense responsive genes induced by transient expression of FOIG_15013 in *N. benthamiana*. The constitutively expressed gene *NbEF1α* was used as internal reference (Situ et al., 2020). Error bars indicate S.D. (n=3). ***, p<0.001.

Hypersensitive response (HR), a form of programmed cell death at the site of pathogen infection, is an important symbol of plant immune activation [32]. To test the effect of FOIG_15013 on host plant immunity, we transiently expressed the milR-87 and its target gene *FOIG_15013* in the leaves of *Nicotiana benthamiana* according to the method described in previous studies [17, 32]. Compared to the GFP control, transient expression of FOIG_15013 caused obvious leaf yellowing in *N. benthamiana* at 48 hours post-agroinfiltration (hpa), while positive control of INF1 caused typical leaf necrosis at the same condition (Fig 9E). And expression of FOIG_15013 could induce significant up-regulation of some pathogenesis-related marker genes including the *PR-1*, *PAL4*, *LOX*, and Osmotin coding gene [33, 34], but not induce up-regulation of HR-related gene the *HIR1* [35] and ethylene response gene the *ERF1* [36]. In contrast, co-expression of milR-87 with FOIG_15013 significantly downregulated these marker genes expression (Fig 9F). The results indicate the FOIG_15013 activates general defense responses in *N. benthamiana*, while milR-87 suppress these defense responses by targeting FOIG_15013 at transcriptional level.

## Discussion

Most fungal genomes encode more than one AGO proteins. The *N. crassa* genome has two, *M. oryzae* three, *C. parasitica* four, and members within the *F. oxysporum* species complex produce two to five AGO proteins [11,14,27]. Eucaryotic AGO proteins are characterized by a bilobal architecture, one contains the N-terminal and PAZ domains, and the orher contains the MID and PIWI domains [37]. The *Foc* genome encodes two AGO proteins, both containing conserved PAZ and PIWI domains. Phylogenetic analysis (Fig 1) showed the FoQDE2 proteins derived from different *formae specialis* were closely clustered in one group, suggesting that FoQDE2 proteins are well conserved and the FoQDE2 is orthologous to QDE2 proteins of *N. crassa*, *F. graminearum*, and *M. oryzae*.

In *F. oxysporum*, several genes related to conidiation were reported [28–31]. Here we selected six of them to verify the function of FoQDE2 in conidial production. qRT-PCR analysis showed that, except for *brlA* and *Htf1*, the transcription of the other four conserved genes (*abaA*, *wetA*, *FoNIIA*, and *StuA*) related to conidiation in *Fusarium* or other fungi was downregulated in Δ*FoQDE2*. The transcript levels of only two of these genes (*FoNIIA* and *StuA*) were restored to the WT level, in the complementation transformant. Previous studies have shown that a nitrite reductase encoding gene, *FoNIIA*, is significantly upregulated during conidiation compared with during vegetative growth in *F. oxysporum*, likely under regulation of a transcription factor Ren1 [28]. *StuA* is involved in the formation of macroconidia in *F. oxysporum* and is also related to spore development and pathogenicity in *F. graminearum* [29, 38]. In our phenotypic analysis, conidiation of Δ*FoQDE2* was markedly lower than that of the WT, suggesting that *FoQDE2* could affect conidial production through downregulating *FoNIIA* and *StuA* expression. In *Aspergillus* and other fungi, three tandem genes (*brlA*, *abaA*, and *wetA*) are considered to be central regulators of conidial production [30, 31]. In this study, the transcript levels of two genes (*abaA* and *wetA*) were increased in the complementation transformant, but were not restored to the WT level, suggesting that the decreased conidiation of Δ*FoQDE2* was not directly related to this central regulatory pathway. And It is worth further identifying QDE2-dependent milRNAs targeting these conidiation-related genes, to further elucidate the regulatory mechanism of *Foc* condiation by miRNAs.

Previous research showed that the AGO protein FgAgo1 and Dicer protein FgDicer2 function in RNAi-mediated gene silencing in *F. graminearum* [39]. Deletion of either of the two genes did not affect the growth and pathogenicity of *F. graminearum* [39]. Our results demonstrate that, compared with WT, the Δ*FoDCL1* and Δ*FoDCL2* mutants have no differences in the tested phenotypic traits, including mycelial growth, conidial production, and sensitivity to H_2_O_2_. The results are consistent with those reported in *F. graminearum* [40, 41], but not the same as in *Va. mali* [42]. In our study, the phenotype of the Δ*FoQDE2* is completely different from that of *F. graminearum*. The differences in pathogenicity of *Foc* and *F. graminearum* may be caused by differences in infection sites. *Foc* infects the host through roots, whereas *F. graminearum* damages the host panicles.

At least five types of milRNA biosynthetic pathways have been reported in fungi. Among them, milR-1 and milR-2 produced by the two biosynthetic pathways, respectively, are dependent on the AGO protein QDE2 in *N. crassa* [11]. Maturation of milR-1 milRNAs, the most abundant milRNAs, requires the AGO protein QDE-2, Dicer, and QIP [23]. In this study, transcript level of milR-87 was significantly lower in Δ*FoQDE2* and Δ*FoDCL2* mutant, compared with that of the WT. The result indicates milR-87 is a FoQDE2-dependent milRNA and belongs to the milR-1 type of milRNA.

We identified a target gene, *FOIG_15013*, of milR-87. *FOIG_15013* encodes a glycosyl hydrolase with an N-terminal signal peptide, therefore we speculate that it may secret out and act as an effector, and likely activates the host defence responses during the infection of *Foc*. Pathogenesis-related proteins (PRs) of plant are usually activated by different biotic and abiotic stresses, including pathogen infection, to confer plant resistance [43, 44]. Examples of PRs include the PR-1 and PAL4, involved in the salicylic acid (SA)-related defense pathways in *N. benthamiana* [44, 45], the Osmotin belonging to PR-5 family and involved in the plant defense against various pathogens [46], and 9-Lipoxygenase (9-LOX) involved in activating local defense against pathogen [34, 47]. In this study, heterologous transient expression of *FOIG_15013* in *N. benthamiana* induced significant up-regulation of these aforementioned resistant marker genes. In contrast, co-expression of milR-87 suppressed the resistant marker genes’ expression. These results support that the *FOIG_15013* encoding glycosyl hydrolase may act as an avirulence factor, thus negatively regulates *Foc* pathogencity, and the milR-87 could suppress the expression of *FOIG_15013* to facilitate infection to the host plant. Future investigation is needed to further elucidate the interaction between FOIG_15013 and plant resistant marker genes expression.

Some milRNAs, found in phytopathogenic fungi *Ve. dahliae* and *Va. mali*, regulate the fungal pathogenicity by targeting their own virulent genes. For example, the VdmilR1, which is independent of Dicer and AGO proteins, regulates fungal virulence at the later stage of inoculation by suppressing a virulence gene (*VdHy1*) expression through increasing histone H3K9 methylation [16]. A Vm-milR37, exclusively expressed in the mycelium, is involved in pathogenicity by targeting a glutathione peroxidase coding gene *VmGP*, which contributes to the oxidative stress response [17]. In the study, a milRNA (milR-87), firstly identified in *Foc*, contributes to the fungal virulence during the early infection stages (24-96 hpi) by targeting glycosyl hydrolase from itself. So it is different from the reported milRNA found in *Ve. dahliae* and *Va. Mali*.

Previous studies showed that aggressive fungal pathogens such as *Botrytis* and *Verticillium* can take up external sRNAs and double-stranded RNAs (dsRNAs) [18, 48]. Applying sRNAs or dsRNAs that target *Botrytis* milRNA synthetic genes *BcDCL1* and *BcDCL2* on the surface of fruits, vegetables, and flowers significantly inhibits gray mold disease [18]. In our study, the virulence of *Foc* to banana leaves was significantly reduced by using exogenous siRNAs, a kind of dsRNAs, to suppress and interfere with the synthesis of milR-87, resulting in reduced disease lesion formation mimicking loss of miR87 function. This result suggest that milR-87 can be used as a target for developing the new and environmentally-friendly fungicides.

In this study, through sRNA sequencing we identify a FoQDE2-dependent milRNA, milR-87, which is significantly upregulated during the early stages of *Foc* infection (24-96 hpi). We further showed that milR-87 promotes fungal virulence through targeting at a glycosyl hydrolase coding gene *FOIG_15013*, which negatively regulates mycelial growth and the virulence of *Foc*. Application of exogenous siRNAs to interfere with the synthesis and expression of milR-87 in *Foc* could effectively suppressed fungal pathogenicity, suggesting that milR-87 could be used as an effective anti-fungal target for developing disease control strategy against banana *Fusarium* wilt.

## Materials and methods

### Fungal strains and culture conditions

The *F. oxysporum* f. sp. *cubense* tropical race 4 strain XJZ2, isolated from Guangdong Province in China [49], was used as the WT for fungal transformation, gene knockout, and milRNA overexpression experiments. All fungal strains were cultured on potato dextrose agar (PDA) for conidiation and mycelial growth. Conidiation was induced as described in our previous study [50]. After a 5-day culture on PDA plates, the colony morphology of the tested strains was recorded. Each experiment was repeated three times independently.

### Oxidative stress test

To test the sensitivity of strains to hydrogen peroxide, a conidial suspension of each strain was spotted on MM or MM supplemented with H_2_O_2_ (2 to 3 mM) and cultured at 28°C. The colony diameter was measured after 4 days’ incubation. A Student’s *t-*test was used to assess significant differences between the WT and mutant.

### Generation of deletion mutants and gene complementation transformants

Deletion mutants of three RNAi-related genes (*FoQDE2*, *FoDCL1*, and *FoDCL2*) in *Foc* were created using a conventional target gene replacement method through homologous recombination. For *FoQDE2* complementation, a 1,713-bp fragment in the promoter region and ORF of *FoQDE2* was cloned into the backbone vector pEX-Zeocin. The Δ*FoQDE2* mutant was transformed with the complementation vector by PEG-mediated transformation as described previously [50].

### Nucleic acid manipulation, small RNA detection and qRT-PCR analysis

Fungal genomic DNA was extracted according to a previously described method [51]. Deletion mutants were verified by PCR and Southern blot analysis according to a previous method [52]. Primers used in the study are shown in Supplementary Table S1. Total RNA was extracted using Trizol reagent (Invitrogen, USA) according to the manufacturer’s protocol. To detect target gene expression, qRT-PCR was carried out as described previously [51]. At least three independent experiments were performed by using three biological replicates.

For detection of small RNAs, a total of 40-60 mg total RNA was separated in a 15% urea-polyacrylamide gel, and transferred to Amersham Hybond^TM^-N^+^ membrane (GE Healthcare, USA). The probes were labelled with biotin using Biotin 3’ End DNA Labeling Kit (Thermo scientific, USA). Hybridization signals were detected by Chemiluminescent Nucleic Acid Detection Module (Thermo scientific, USA). Signal intensity was quantified using Image Lab 6.1.0 software (BIO-RAD, USA).

An All-in-One miRNA qRT-PCR Detection Kit (GeneCopoeia, Rockville, MD) was used for milRNA expression analysis [12]. Briefly, a PolyA tail was added to total RNAs using PolyA polymerase (NEB, USA). The RNA was then reverse-transcribed with an oligo-dT adaptor. MilRNA expression was detected using qRT-PCR, as described above, except that the reference gene was replaced by snRNA U4 in *Foc*. At least three biological replicates for each sample were performed.

### Small RNA library construction and sequencing

Total RNAs of the WT strain XJZ2 and *FoQDE2* deletion mutant Δ*FoQDE2* were extracted as described above. RNA integrity was assessed and quantified with a Bioanalyzer 2100 (Agilent, USA) and only qualified RNA samples (RNA integrity number, RIN: 7–10) were used for sRNA library construction. Three different samples of every strain were used for sRNA library preparation and sRNA sequencing. Small RNA libraries were prepared using NEB Next^®^ Small RNA Library Prep Set for Illumina (NEB, USA) according to the manufacturer’s protocol. Briefly, total RNAs were reverse transcribed and indexed. Then, cDNAs were separated on a 6% polyacrylamide gel and cDNAs ranging from between 140 and 160 nt, corresponding to 20–40 nt sRNAs, were cut and recycled. The size and concentration of the sRNA sequencing libraries were assessed again using Bioanalyzer 2100. Three sRNA libraries from the WT fungus and two from the Δ*FoQDE2* mutant were qualified for sRNA sequencing. Sequencing was performed on an Illumina MiSeq platform at the University of Massachusetts, Amherst using an MiSeq Reagent Kit v2 for single-end reads at a length of 50 bp.

### Small RNA data analysis and milRNA prediction

Raw sequencing data were trimmed and analyzed using CLC Genomics workbench software v10.1 (CLC bio) and only reads with a read count number larger than 10 and length between 16 and 40 nt were kept for further analysis. The *Foc* II5 (TR4 strain) genome released by the Broad Institute was used as a reference sequence. To examine the origin of the sRNA, reads mapped to the *Foc*II5 genome were collected and sequentially mapped to the rRNA, tRNA, snRNA, snoRNA, repeat, exonic, intronic, and 3′ and 5′ UTRs regions of coding genes, and intergenic regions (the region 1000 nt upstream and 1000 nt downstream of genes). To reduce false positive sRNAs induced by mRNA degradation, only reads mapped to the UTR, intron, and intergenic regions were used for sRNA length distribution and sRNA locus identification. Overlapping and adjacent reads were grouped and only sequences with a consensus length of less than 300 nt were selected and uploaded to the RNAfold web server (http://rna.tbi.univie.ac.at//cgi-bin/RNAWebSuite/RNAfold.cgi) for RNA secondary structure prediction.

To identify FoQDE2-dependent sRNAs and their loci, reads from the WT and Δ*FoQDE2* were compared to each other. Read counts were normalized by calculating their transcripts per million (TPM) value. After normalization, log2 ratios were calculated using the TPM value from two Δ*FoQDE2* mutants and three WT XJZ2 strains. Only sRNAs with log2 ratios of less than –1 or greater than 1 and a *p* value of less than 0.05 were considered as significantly expressed. The sRNAs, which were significantly downregulated in Δ*FoQDE2* and their precursors with a typical secondary stem and loop structure were considered as FoQDE2-dependent milRNAs. For all predicted and differentially expressed milRNAs, transcript levels were verified by qRT-PCR analysis.

### Generation of the milRNA deletion mutants and overexpression transformants

The milRNA deletion mutants were created by replacing the precursor (about 300 bp) of milRNA through homologous recombination as functional gene deletion we previously described [50]. The vector pSilent-1 was donated by Dr. Hitoshi Nakayashiki for milRNA overexpression. The milRNA precursor was amplified from the *Foc* genome and then cloned into the pSilent-1 vector through digestion with *Hin*dIII and *Kpn*I. The inserted fragment was verified by sequencing and the correct vector was used for transformation of the WT strain XJZ2 of *Foc*. Transformants were identified by PCR with primers given in the table S1.

### Transfection of siRNAs and inhibitor to suppress the production of milR-87 in *Foc*

Protoplast preparation was performed as described previously [50]. Double-strand siRNAs (siRNA-1 and siRNA-2) and single-strand inhibitor against the precursor sequence of milR-87, as well as a random sRNA sequence (24 nt) serving as a negative control (NC) were designed and synthesized by General Biosystems company. According to the protocol provided by transfection reagent, a total of 100 nM siRNA oligos or inhibitor was transfected into the prepared protoplast of *Foc* using lipofectamine 2000 (Invitrogen). After two days culture, conidia suspensions of the transfected strains and positive control of the ΔmilR-87 mutant, as well as non-transfected strain of CK were inoculated on the banana leaves. After three days, the lesions on banana leaves were recorded and measured for accessing virulence of different treatments. The total RNAs of transfected strains cultured at 28°C for 7 days were extracted for milRNA detection.

### Fungal invasion assays and pathogenicity test

The ability of mycelia to penetrate cellophane and the invasive growth on tomato fruit surfaces were compared between WT and Δ*FoQDE2* according to a previous description [50]. The pathogenicity on banana seedlings was assessed as described previously [4]. A total of 30 banana plantlets were used for each treatment. The severity of internal symptoms was recorded with disease indexes and a Student’s *t*-test was used to assess significant differences.

### MilRNA target gene identification and transient expression in ***Nicotiana benthamiana***

Based on the sequence of milR-87, its target genes in *Foc* genome were predicted using psRNATarget online software. In which the genes significantly up-regulated in the mutants ΔmilR-87 and Δ*FoQDE2* were screened by qRT-PCR and selected for target gene identification. According to the target site of the milRNA, a GFP marker gene was fused to the 3 ’terminal of the candidate target gene. And a point-mutation of the milR-87 targeted site was introduced into the WT. The GFP fluorescence intensity representing the expression of the target gene was accessed by confocal microscopy. Furthermore, the expression of GFP and target genes was quantified by qRT-PCR.

Transient expression vector was constructed according to the method described by [32]. Briefly, precursor of the milR-87 and *FOIG_15013* were introduced to vector pBINGFP. The recombinant vectors were transformed into *Agrobacterium tumefaciens* GV3101. For transient expression, transformed *A. tumefaciens* cultures were injected into *N. benthamiana* leaves [13]. After 48 h, the injected leaves were photographed and then harvested for detecting mRNA and protein levels of the FOIG_15013, as well as transcript level of resistant marker genes described previously in *N. benthamiana* [34,35,44–46].

## Supporting information

S1 Table. Primers used in this study.

S2 Table. Daily colony diameters of tested isolates cultured on PDA plates.

S1 Fig. Verification of the *FoQDE2* deletion and complemented mutants by PCR and Southern blot analysis. (**A**) Schematic diagram of *FoQDE2* gene deletion and complementation.

Short arrows in the figure show the primer sites in the study. (**B**) PCR Identification of the *FoQDE2* deletion mutants. M, DL5000 DNA ladder purchased from TAKARA; 1-6, the different *FoQDE2* gene deletion mutants; WT, wild type strain XJZ2 of *Foc*; ddH_2_O, negative control. (**C**) Southern blot analysis with probes of the Hygromycin resistance gene fragment and the respective left border fragments of *FoQDE2*, *FoDCL1* and *FoDCL2*. WT, indicates wild type strain XJZ2 of *Foc*; 1-6, the different *FoQDE2* gene deletion mutants; 7-10, the different *FoDCL1* gene deletion mutants; 11-14, the different *FoDCL2* gene deletion mutants. Genomic DNA was digested by *Hind* III overnight, separated in a 0.8% agarose gel, blotted onto a N+ nylon membrane, and hybridized with the Dig-labeled HYG probe amplified with the primer pair HYG-F/HYG-R and the LB probe amplified with the primer pair P1/LB-R. (**D**) PCR Identification of the *FoQDE2* complemented transformants. M, DL2000 DNA ladder purchased from TAKARA; 1-4, the different *FoQDE2* complemented transformants; 5, the WT strain; 6, the *FoQDE2* gene deletion mutant; 7, complimentary vector DNA as positive control; 8, ddH_2_O as negative control.

**S2 Fig. Examination of relative transcript levels of small RNA biosynthesis genes.** (**A**) Relative transcript levels of *FoQDE2*, *FoDCL1*, and *FoDCL2* were examined by quantitative real-time PCR (qRT-PCR) analysis in the WT strain and their corresponding deletion mutants. (B) Relative transcript levels of *FoQDE2* in the WT strain, *FoQDE2* deletion mutant and *FoQDE2* complemented transformants.

**S3 Fig. Verification of the milR-87 deletion and overexpression mutants by PCR, qRT-PCR and Northern blot.** (**A**) Schematic diagram for deletion and overexpression of milRNA (milR-87) in *Fusarium oxysporum* f. sp. *cubense* (*Foc*). (**B**) PCR Identification of the milR-87 deletion mutants. M1 is DL5000 DNA ladder. (**C**) PCR Identification of the milR-87 overexpression mutants. M2 is DL2000 DNA ladder. (**D**) qRT-PCR detection of milR-87 in different mutants of *Foc*. (**E**) Northern blot analysis of milR-87 in the different mutants of *Foc*.

**S4 Fig. Identification of milR-87 target gene in *Fusarium oxysporum* f. sp. *cubense* (*Foc*).** Relative transcript levels of the different target genes predicted online were examined by qRT-PCR in the WT strain XJZ2, the ΔmilR-87 mutant and the Δ*FoQDE2* mutant of *Foc*.

**S5 Fig. PCR identification of the *FOIG_15013* deletion mutants of *Fusarium oxysporum* f. sp. *cubense* (*Foc*).** (**A**) PCR Identification of the *FOIG_15013* deletion mutants. M, DL5000 DNA ladder purchased from TAKARA. Δ*FOIG_15013* -9/-10/-19, the different *FOIG_15013* gene deletion mutants; XJZ2, the wild type strain of *Foc*. ddH_2_O, negative control. (**B**) qRT-PCR detection of *FOIG_15013* in the different Δ*FOIG_15013* mutants and the wild type strain of *Foc*.

## Acknowledgements

We thank Professor Yi-zhen Deng of South China Agricultural University for critical suggestions on the manuscript writing. We are grateful to Professor Button B. Yang of Toronto University for advices on siRNA experiments.

## Author Contributions

**Conceptualization:** Minhui Li, Li-jun Ma, Zide Jiang.

**Data curation:** Minhui Li, Lifei Xie, Meng Wang, Yong Zhang, Jin Zeng.

**Formal analysis:** Minhui Li, Huaping Li, Zide Jiang.

**Funding acquisition:** Minhui Li, Huaping Li.

**Investigation:** Yong Zhang, Guanghui Kong, Pingen Xi.

**Methodology:** Lifei Xie, Meng Wang, Jiaqi Zhong, Jin Zeng, Yong Zhang, Li-jun Ma.

**Project administration:** Minhui Li, Li-jun Ma, Zide Jiang.

**Writing – original draft:** Minhui Li, Yilian Lin

**Writing – review & editing:** Li-jun Ma, Zide Jiang.

## Notes

### Competing Interest Statement

The authors have declared no competing interest.

